# HyINDEL – A Hybrid approach for Detection of Insertions and Deletions

**DOI:** 10.1101/2021.10.08.463662

**Authors:** Alok Thatikunta, Nita Parekh

## Abstract

Insertion and deletion (INDELs) mutations, the most common type of structural variation in the human genome, have been implicated in numerous human traits and diseases including rare genetic disorders and cancer. Next generation sequencing (NGS) technologies have drastically reduced the cost of sequencing whole genomes, greatly contributing to genome-wide detection of structural variants. However, due to large variations in INDEL sizes and presence of low complexity and repeat regions, their detection remains a challenge. Here we present a hybrid approach, HyINDEL, which integrates clustering, split-mapping and assembly-based approaches, for the detection of INDELs of all sizes (from small to large) and also identifies the insertion sequences. The method starts with identifying clusters of discordant and soft-clip reads which are validated by depth-of-coverage and alignment of soft-clip reads to identify candidate INDELs, while the assembly -based approach is used in identifying the insertion sequence. Performance of HyINDEL is evaluated on both simulated and real datasets and compared with state-of-the-art tools. A significant improvement in recall and F-score metrics as well as in breakpoint support is observed on using soft-clip alignments. It is freely available at https://github.com/alok123t/HyINDEL.

## I. Introduction

Insertion and deletion (INDELs) mutations refer to contiguous stretch of nucleotides inserted or deleted in genomic DNA and are one of the most common class of mutations in human genome [1]. These can affect large number of bases and lead to gain/loss of functional part of genes, change in gene expression levels, frame-shift errors and integration of foreign DNA into the genome. Their detection has important clinical applications as these are implicated in many genetic, neural and oncologic diseases [2]. In cancer, INDELs are a common mechanism of kinase activation. Kinase inhibitors being an important class of targeted cancer therapeutics, detection of INDELs plays an important role in cancer genomics.

Various methods have been proposed for detection of insertions and deletions in next generation sequencing data. Discordant-pairs (abnormal insert size or orientation), split read (chimeric alignment of reads) and read-depth (abnormal coverage) are various signals identified to be associated with INDELs. The method for INDEL detection are classified into four categories based on the signals used: Clustering approach (e.g., BreakDancer [3], PEMer [4]), Split-mapping approach (e.g., CREST [5], Socrates [6]), Assembly approach (e.g., NovelSeq [7], Pamir [8]), and Hybrid approach (e.g., LUMPY [9], SoftSV [10]). Since a single signal cannot comprehensively detect all types and sizes of INDELs, it is desirable to incorporate multiple signals for accurate detection. With this objective, we propose a hybrid approach, HyINDEL, that integrates information of discordant and split reads, as well as partially mapped split reads (soft-clip reads) and depth of coverage to identify and validate candidate INDELs. Using the assembly-based approach, soft-clipped reads are aligned to construct the entire insertion sequence. It is implemented in C++ using Bamtools [11] and is available with open source MIT license at https://github.com/alok123t/HyINDEL.

## II. Materials and Methods

Below we discuss the proposed approach, describing key idea of using soft-clip reads and depth of coverage. First, we discuss the distribution of different types of reads observed in the vicinity of insertions and deletions which help in identifying probable INDEL sites.

### Distribution of Reads at a Deletion Locus

A contiguous sequence of bases absent in sample with respect to reference is known as a deletion. It is represented by two breakpoints on the reference, referred to as 5′ and 3′ breakpoints, the sequence between which is missing in the sample (**Fig. 1**(a)). It may be noted that paired-end reads in the vicinity of a deletion event can be categorized into different types based on the orientation and insert size, when mapped to reference genome. A paired-end read that does not cross INDEL breakpoint, aligns to the same chromosome of reference with same inward orientation and insert size within the range (span_min_, span_max_), where span_min_ = I_median_ - *k*δ and span_max_ = I_median_ + *k*δ, I_median_ and δ are median and standard deviation of the insert size, (default *k*=3). These are called *concordant* reads (reads 1 and 8 in **Fig. 1**(a)). Anomalous paired-end reads are those that cross one of the deletion breakpoints. In this case two reads of a pair do not align concordantly (i.e., insert size > span_max_) and are called *discordant* reads (reads 3, 4, 5 and 6 in **Fig. 1**(a)). When a part of the read aligns at 5′ breakpoint and the other part aligns at 3′ breakpoint of the deletion (or vice-versa), because of the split nature of alignment these are called *split* reads (reads 3 and 6 in **Fig. 1**(a)). If only one part of the read aligns to the reference while the other part is unmapped, then such reads are called *soft-clip* reads. Reads marked with soft-clip at 3′ end of alignment (e.g., 70M30S in CIGAR string) provide information of 5′ breakpoint of the deletion (read 2 in **Fig. 1**(a)), while those marked with a soft-clip at 5′ end of alignment (e.g., 20S80M) provide information about 3′ breakpoint (read 7 in **Fig. 1**(a)). Thus, in the vicinity of a deletion event, cluster of discordant, soft-clipped and split reads are observed which are helpful in accurately detecting breakpoints of the deletion region.

**Fig. 1.**
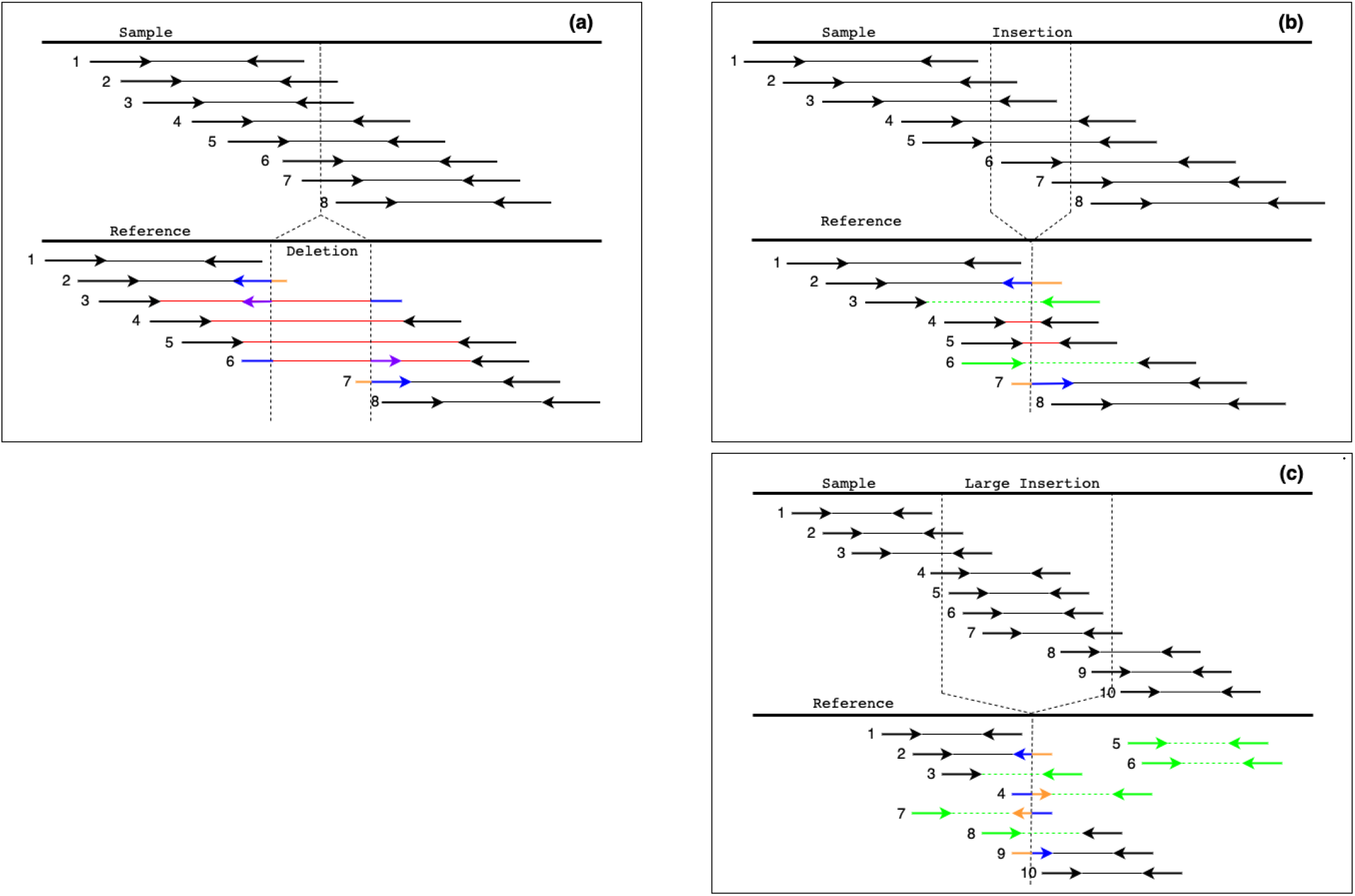
(a) A deletion event shown. Paired-end reads (1 and 8) shown in black, are concordant reads that correctly align to reference genome, while large insert size is observed in case of discordant reads (marked in red). Partial ends of reads (shown in blue for reads 2 and 7) denote the aligned part, while unaligned parts (shown in orange) are the soft-clipped part of the reads. Reads showed in purple (reads 3 and 6) are split reads, wherein both the partial ends of the read are aligned to reference. (b) An insertion event shown. Here, black paired-end reads (1 and 8) are concordant reads that correctly align to reference genome, while small insert size is observed in case of discordant reads (reads 4 and 5 in red). Partially aligned reads (reads 2 and 7 shown in blue) have their soft-clipped part (shown in orange) unaligned. Reads 3 and 6 marked in green denote unaligned end, called one-end anchored (OEA) reads. (c) A large insertion event is shown. Paired-end reads (1 and 10) shown in black are concordant reads. Partial ends of reads shown in blue denote the aligned part, while unaligned part (shown in orange) are the soft-clipped part of the reads 2, 4, 7 and 9. Reads for which only end is mapped, called one-end anchored reads (reads 3 and 8) and reads for which neither end of the paired-end reads are aligned (reads 5 and 6 shown in green), are called orphan reads.

### Distribution of Reads at an Insertion Locus

A continuous sequence of bases present in sample but missing in reference as shown in **Fig. 1(b)** corresponds to an insertion event. It is represented by a single breakpoint on the reference. As in the case of deletion, paired-end reads in the vicinity of an insertion event can be categorized into different types based on orientation and insert size when mapped to reference genome. Paired-end reads that do not cross the insertion breakpoint, aligns to same chromosome of reference with same inward orientation and insert size are called *concordant* reads (reads 1 and 8 in **Fig. 1(b)**). Paired-end reads with a short insert size (< span_min_) but with same inward orientation are called *discordant* reads (reads 4 and 5 in **Fig. 1(b)**). When one of the read of a pair spans the insertion breakpoint (reads 2 and 7 in **Fig. 1(b)**), 5′ (3′) end of the read is partially aligned, called *soft-clip* reads. These help in precisely detecting the breakpoint and prefix (suffix) of the ‘insertion’ sequence. If the insert size is larger than read length, typically only one read of the pair is mapped to reference genome (reads 3 and 6 in **Fig. 1(b)**) and are called *one-end anchored* (OEA) reads. For identifying insertions larger than the insert size of the library, in addition to soft-clip (reads 2, 4, 7 and 9) and one-end anchor reads (reads 3 and 8), orphan reads (i.e., neither ends of a paired-end read map the reference genome) are also considered (reads 5 and 6 in **Fig. 1 (c)**). The proposed approach uses information from all these different types of reads to detect the location of insertion breakpoint and construct the insertion sequence.

#### A. Pre-Processing

Eukaryotic genomes are rich in repeat sequences and low-complexity regions, especially the centromeres and telomeres. Reads originating from these regions ambiguously align to multiple regions in reference genome. Consequently, several discordant and soft-clip reads with abnormally high depth of coverage are observed in these regions. To avoid predicting spurious variants in these regions, read depth profile of sample genome is constructed in non-overlapping bins of size 1000bp. The median coverage (c_median_) of sample is computed and bins having read depth (> 3 × c_median_) are discarded to filter low complexity regions. This step is performed using Mosdepth [12] in our pipeline. Also, alignments with low mapping quality (< 20) are not considered for variant detection by parsing the BAM file.

#### B. Detection of Deletions

##### Clustering Discordant reads

As seen in **Fig. 1(a)**, the discordant reads in this case exhibit larger insert size, but same inward orientation. Let 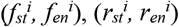 denote start and end of the alignment of forward and reverse reads of *i*^*th*^ discordant paired-end read on the reference. Two paired-end discordant reads (say, reads 3 and 4 in **Fig. 1(a)**) are merged into a single cluster if the reciprocal overlap of the region between *f*_*st*_ and *r*_*en*_ for the two reads is ≥ 0.65 (i.e., *f*_*st*_ ^*4*^ to *r*_*en*_ ^*3*^ region is ≥ 0.65). Other paired-end reads satisfying this criterion with any of the reads of this cluster are merged into this cluster. This results in clusters of discordant paired-end reads, which indicate approximate location of breakpoints of probable deletion events. We expect 5′ breakpoint of the deletion to lie within the interval (*f*_*en*_^*i*^, *f*_*en*_^*i*^ *+ span*_*max*_) and 3′ breakpoint within (*r*_*st*_^*j*^ *-span*_*max*_, *r*_*stj*_), where *i* and *j* correspond to reads with farthest 5′ and 3′ coordinates in the cluster, respectively.

##### Clustering Soft-clip reads

Two soft-clipped reads support the 5′ (3′) breakpoint of the same deletion event if start (end) of soft-clipped part are within 5bp and exhibit high sequence similarity over the overlap region (≥ 90%), and belong to the same cluster. The sequence similarity is assessed by carrying out semi-global alignment using the scoring scheme (1, −1, −1) for match, mismatch and gap penalty respectively. Soft-clip cluster containing reads with soft-clip at 5′ (3′) end is known as downstream (upstream) soft-clip cluster respectively. The soft-clip clusters thus constructed are used in the detection of breakpoint. A split-read that has an overlap of ≥ 90% with any read of 5′ (3′) softclip cluster, is merged with the respective soft-clip cluster.

By analyzing the discordant clusters, approximate location of a deletion region is obtained. On identifying the upstream and downstream soft-clip clusters that overlap with the discordant clusters, the precise location of the breakpoints can be obtained. Any discordant/soft-clip cluster having less than c_median_/10 reads are discarded, where c_median_ is the median depth-of-coverage of sample. Below we briefly discuss our approach in the detection of small (50, 500) and large deletions (> 500).

##### Detection of Large deletions

Each discordant cluster provides approximate location of 5′ and 3′ breakpoints of a deletion event within the interval 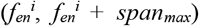 and 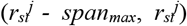, where *i* and *j* correspond to reads with farthest 5′ and 3′ coordinates in the cluster, respectively. If a soft-clip cluster has its coordinates lying within the interval 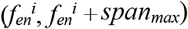, it defines the 5′ breakpoint, and the soft-clip cluster with coordinates lying within the interval 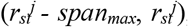 defines the 3′ breakpoint region. Two soft-clip clusters define the same deletion event if alignment of reads from the two clusters exhibit high sequence similarity (as discussed below). If no soft-clip cluster pair is found to map the discordant cluster, the candidate deletion is reported as *imprecise* deletion event.

##### Detection of Small deletions

Small deletions (< *L*_*small*_ = *k*δ, *k* = 3 and δ corresponds to standard deviation of the insert size) are missed by the above approach as in this case all paired-end reads around the breakpoint get classified as concordant reads. Thus, only soft-clip clusters are available for the detection of small deletions. In this case for each upstream soft-clip cluster, a downstream soft-clip cluster within a distance of *L*_*small*_ and exhibiting high sequence similarity between inter-cluster reads is identified, indicating they represent the same deletion event (as discussed below).

##### Confirmation by alignment of soft-clipped reads

As seen above, each deletion event is represented by an upstream and a downstream soft-clip cluster. For correctly paired upstream and downstream soft-clip clusters that define the same deletion event, unaligned regions of soft-clip reads from the upstream cluster would be similar to the mapped regions of soft-clip reads from the downstream cluster and vice-versa, as shown in **Fig. 1**(a). Hence, a semi-global alignment of reads from the upstream soft-clip cluster and the corresponding downstream cluster is performed. Two clusters are considered to be associated to the same deletion event if average inter-alignment score is ≥ ½ × *r*_*len*_, where *r*_*len*_ denotes read length. Three largest soft-clip reads (based on soft-clipped regions) from each cluster are considered for alignment since they are expected to have the highest overlapping regions. This alignment step is skipped, if the two clusters contain 3 or more split-reads with their 5′ and 3′ ends mapping respectively to the 5′ and 3′ soft-clip clusters. The deletion breakpoints are then identified as the median start (end) location of soft-clip position of upstream (downstream) cluster respectively.

#### C. Detection of Insertions

As is clear from **Fig. 1**(b), each insertion event is characterized by a cluster of soft-clip reads upstream and downstream of the breakpoint (e.g., reads 2 and 7). For correctly paired upstream and downstream soft-clip clusters that define the same insertion event, we expect the start of soft-clip position from upstream cluster to be the same as end of soft-clip position from the downstream cluster and represent candidate insertion site. For each such site, OEA reads within ± span_max_ of the breakpoint are extracted. To identify the insertion sequence itself, *de novo* assembly of partially aligned soft-clip reads, unaligned ends of OEAs and orphan reads is performed using Minia [13], a short-read assembler based on a de-Bruijn graph. This results in a set of contigs (in fasta format) corresponding to each insertion event. The contig is expected to span the entire insertion sequence and also partially contain region adjacent to the insertion breakpoint. When this contig is aligned to the reference, we expect a split alignment from which we can identify the insertion breakpoint and sequence as described below. Alignment to reference is done using Minimap2 [14], a sequence alignment program for aligning long reads or assemblies to a reference genome.

##### Identification of insertion breakpoint and sequence

The assembled contig comprises a prefix, insertion sequence and suffix, with both prefix and suffix ends mapping to the reference. Small insertions are directly represented as CIGAR string in the alignment file (e.g., 40M120I50M). The CIGAR string is parsed to obtain the position of insertion with respect to alignment position of the read. Based on the position and length of insertion from the CIGAR string, the insertion sequence is identified.

In case of large insertions, two split-alignments of the same contig with reference is observed. This is because the assembled contig contains insertion sequence which is much larger than the read length and two flanking regions that map to reference. Thus, in one of the split alignments, prefix of the contig is aligned to reference, and the remaining part of contig is marked with a soft-clip, while in the other, suffix of the contig is aligned to reference and the remaining part marked as soft-clip. The start position of soft-clip is identified as the insertion position. The soft-clip sequence (after excluding the other split-aligned portion of contig) is identified as insertion sequence. In case split-reads are not available, the insertion event is reported as *imprecise* insertion event with a partial insertion sequence, as in this case we are unable to completely assemble the insertion sequence into a single contig.

#### D. Post-processing

INDEL events that have low support are filtered out. For large deletions identified using both discordant and soft-clip signals, minimum support of reads required is > threshold = c_median_/3. For imprecise and small deletions identified using only soft-clip reads, a threshold of c_median_/6 is used. In the case of homozygous deletions, we do not expect any reads present in the deletion region, while in case of heterozygous deletions, approximately half of the reads are expected to be present. Hence, we compute the ratio of sequence coverage of candidate deletion region (cov_event_) to that of its upstream and downstream flanking regions (cov_flank_) of size 1000bp each. For a candidate deletion, if the ratio cov_event_/cov_flank_ < 0.2 (for both flanks), it is referred to as a homozygous deletion, and if it lies in the interval 0.2 ≤ cov_event_/cov_flank_ ≤ 0.9, it is reported as a heterozygous deletion, remaining events are classified as complex variants. The variants thus predicted are reported in VCF output format (v4.2).

## III. Results

Performance of the proposed approach is evaluated on both simulated and real data using measures Precision, Recall and F-score, given by equations (1)-(3). Precision is defined as the ratio of true positive events to total number of events predicted, while recall is defined as the ratio of true positive events to total number of actual events. F-score is computed as the harmonic mean of these two measures, Precision and Recall. A deletion is considered as true positive if the reciprocal overlap of predicted and actual deletion region is at least 50%. An insertion is considered true positive if the distance between predicted and actual insertion site is within 10bp.

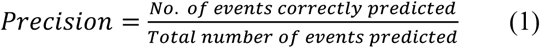

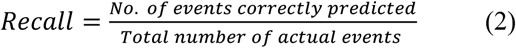

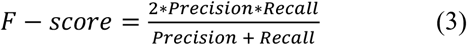

### A. Simulated Experiment

We first evaluate the performance of HyINDEL on simulated data. For this, a diploid sample is constructed by inserting variants into the human genome (assembly GRCh37). Location of insertions and deletions are identified randomly using SVSim [17] and respectively saved in two lists. Each list consists of 750 entries with 375 each from the two size ranges: small (50, 500) and large (500, 10000). First haploid sample is constructed by inserting 375 insertions and 375 deletions in reference genome. The second haploid sample is constructed by inserting all the 750 insertions and 750 deletions in reference genome. The two haploid samples are then merged to obtain a diploid sample containing 750 homozygous (375 insertions, 375 deletions) and 750 heterozygous (375 insertions, 375 deletions) variants. Paired-end reads of length 100bp are generated using ART simulator [19] to simulate reads from Illumina HiSeq 2500. Mean and standard deviation of the insert size for the paired-end reads is set to 350bp and 20bp respectively. Three sets of reads corresponding to sequencing coverage 10×, 20× and 30× are generated. Reads are aligned using BWA-MEM [18] to human genome (assembly GRCh37). The resulting BAM files are sorted by coordinate and indexed using Samtools [20].

Precision, Recall and F-score measures are summarized for deletions (Table-I) and insertions (Table-II) for both homozygous and heterozygous variants at varying sequence coverages. Recall and F-score values are observed to be higher for deletions compared to insertions and improve with increase in sequencing coverage. As expected, recall values are higher for detection of homozygous events compared to heterozygous INDEL events. A high precision value of ~1 observed for all cases clearly indicate the reliability of our predictions. The INDEL events missed by our approach are mainly from regions with low support (and filtered out).

**TABLE I.**
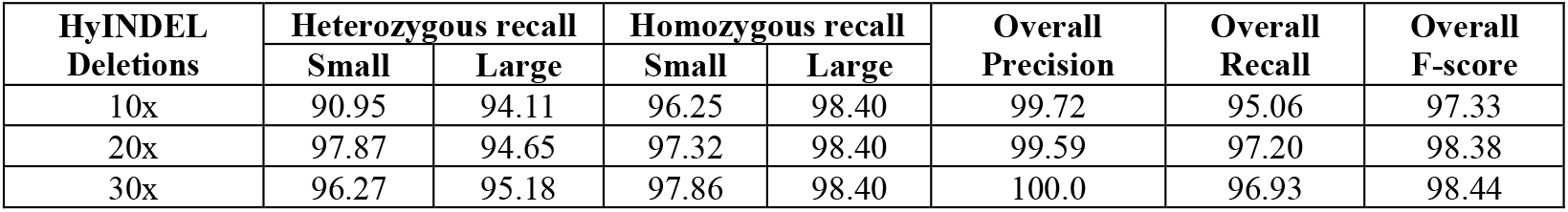
Precision, Recall and F-score metrics for predicting deletions using HyINDEL on simulated data at varying sequence coverages

**TABLE II.**
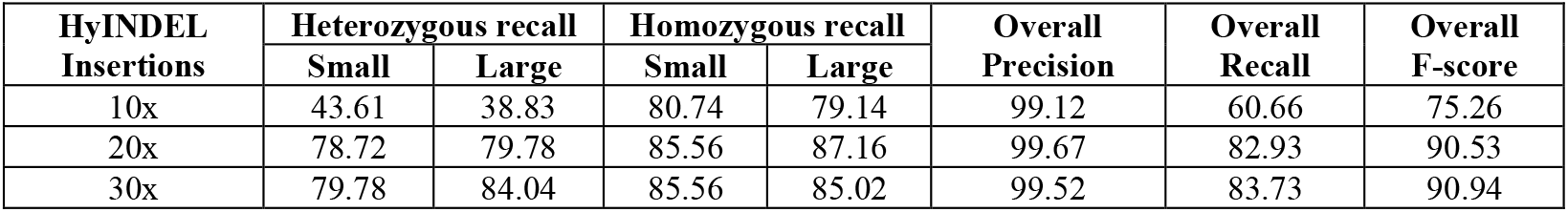
Precision, Recall and F-score metrics for predicting insertions using HyINDEL, on simulated data at varying sequence coverage

| HyINDEL<br>Insertions | Heterozygous recall |  | Homozygous recall |  | Overall<br>Precision | Overall<br>Recall | Overall<br>F-score |
| --- | --- | --- | --- | --- | --- | --- | --- |
|  | Small | Large | Small | Large |  |  |  |
| 10x | 43.61 | 38.83 | 80.74 | 79.14 | 99.12 | 60.66 | 75.26 |
| 20x | 78.72 | 79.78 | 85.56 | 87.16 | 99.67 | 82.93 | 90.53 |
| 30x | 79.78 | 84.04 | 85.56 | 85.02 | 99.52 | 83.73 | 90.94 |

Next, the predictions from HyINDEL are compared with other tools on simulated data, using the metrics, Precision, Recall and F-score at different sequence coverage. The results are summarized in Tables III and IV for deletions and insertions respectively. It may be noted that precision value of HyINDEL is higher or comparable with other tools for all cases indicating the reliability of our predictions. Also, Recall and F-score values are comparable (or marginally higher) to those of other tools on simulated data. Majority of insertion events missed by all the tools correspond to heterozygous insertions, due to low read support.

**TABLE III.**
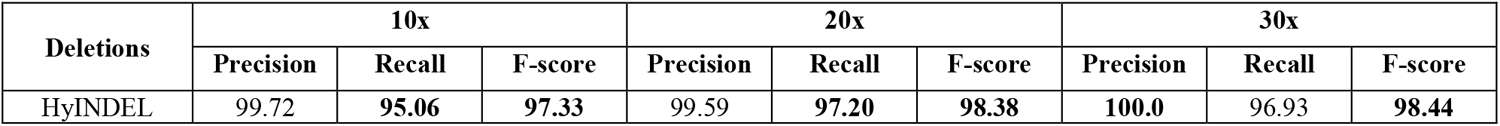
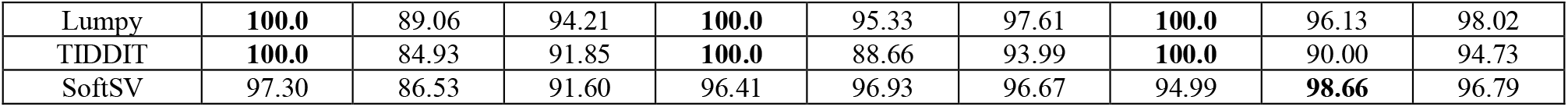
Precision, Recall and F-score metrics for predicting deletions using HyINDEL with other tools on simulated data at varying sequence coverage

**TABLE IV.** Precision, Recall and F-score metrics for predicting insertions using HyINDEL with other tools on simulated data at varying sequence coverage

| Insertions | 10x |  |  | 20x |  |  | 30x |  |  |
| --- | --- | --- | --- | --- | --- | --- | --- | --- | --- |
|  | Precision | Recall | F-score | Precision | Recall | F-score | Precision | Recall | F-score |
| HyINDEL | **99.12** | 60.66 | 75.26 | **99.67** | **82.93** | **90.53** | **99.52** | 83.73 | 90.94 |
| Pamir | 96.88 | 37.33 | 53.89 | 97.64 | 71.86 | 82.79 | 95.73 | **86.80** | **91.04** |
| Popins | 88.12 | **66.26** | **75.64** | 98.98 | 78.00 | 87.24 | 99.68 | 83.60 | 90.93 |

The error in breakpoint predictions for an INDEL event is defined as the absolute difference between true and predicted locations. The breakpoint error for HyINDEL is compared with other tools for the detection of deletions and insertions in **Fig. 2 (a)** and **(b)** respectively. It is observed to be lower for HyINDEL (median breakpoint error 1bp at 30x sequence coverage) possibly because the number of reads supporting the breakpoint is larger in this case compared to other tools, as seen in **Fig. 3** for the detection of deletions. This is because of using soft-clip reads along with split reads in our method. However, the number of outliers observed in SoftSV is least compared to all other tools is because it does not report imprecise events.

**Fig 2.**
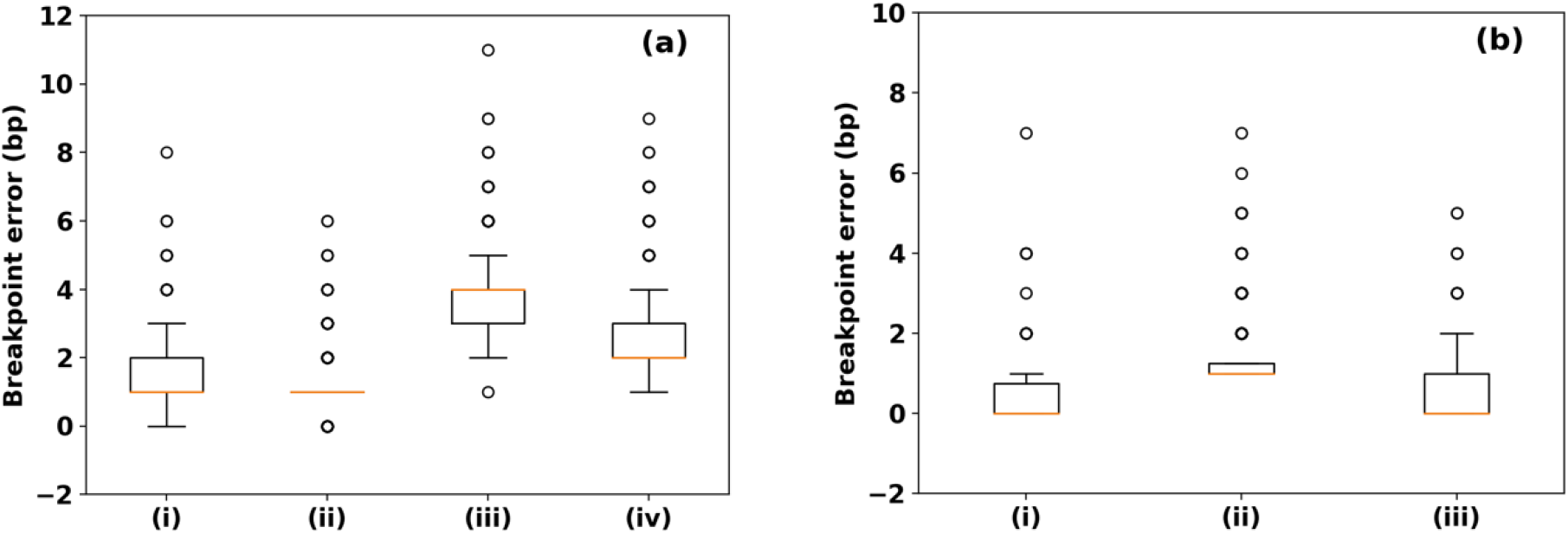
Breakpoint error for simulated data at 30x sequencing coverage in detection of (a) deletions using (i) HyINDEL, (ii) Lumpy, (iii) TIDDIT and (iv) SoftSV and (b) insertions using (i) HyINDEL (ii) Pamir (iii) Popins.

**Fig 3.**
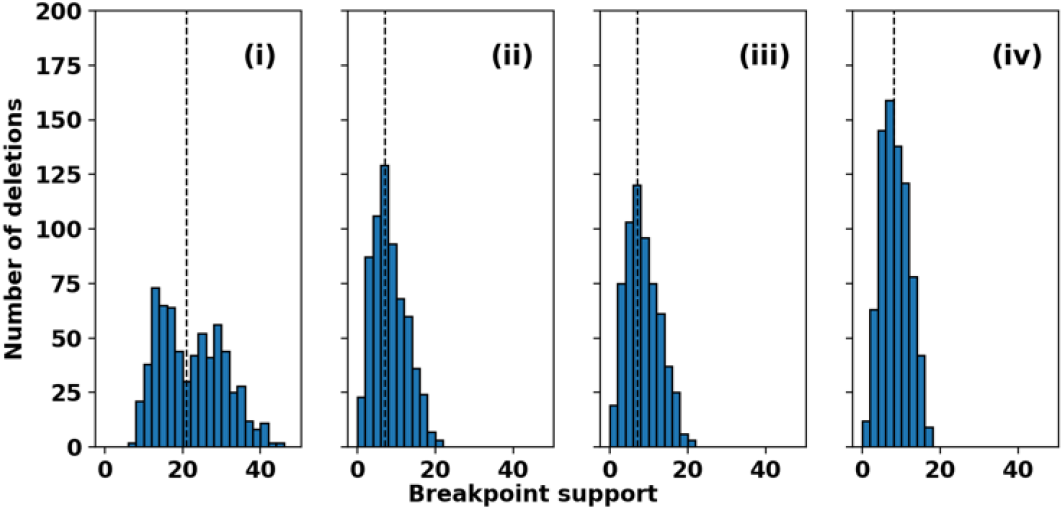
Breakpoint support for deletions (without discordant reads) shown on simulated data at 30x sequencing coverage: (i) HyINDEL, (ii) Lumpy, (iii) TIDDIT and (iv) SoftSV.

### B. Real dataset

Next, for evaluating the performance of our method on real dataset, the most widely studied NA12878 sample is considered. Reads are downloaded from Illumina Platinum genomes repository (ENA accession: PRJEB3381) [21]. It contains paired-end reads of length 101bp generated using Illumina HiSeq 2000. The reads were aligned using BWA-MEM [11] to human genome (assembly GRCh37). The resulting BAM files are sorted by coordinate and then indexed using Samtools [12]. The sorted BAM file and its index are given as input to our tool. The median insert size is estimated to be 318 with standard deviation 78 using Picard tools [22]. The average genome coverage was estimated to be 52× using Mosdepth [12].

Annotations for INDEL events for NA12878 sample is available from two resources, *viz*., Genome in a bottle (GIAB) [23] and Database of Genomic Variants (DGV) [24]. For evaluation, INDELs of size ≥ 50bp are considered. For deletions, a prediction is considered true positive if there is at least a reciprocal overlap of 50% between the actual and predicted event. For insertions, a prediction is considered true positive if it lies within 200bp of an actual event. The results are summarized in Table V and VI for deletions, and in Table-VII for insertions. Performance of HyINDEL is compared with the five tools, for annotations from both GIAB and DGV.

**TABLE V.**
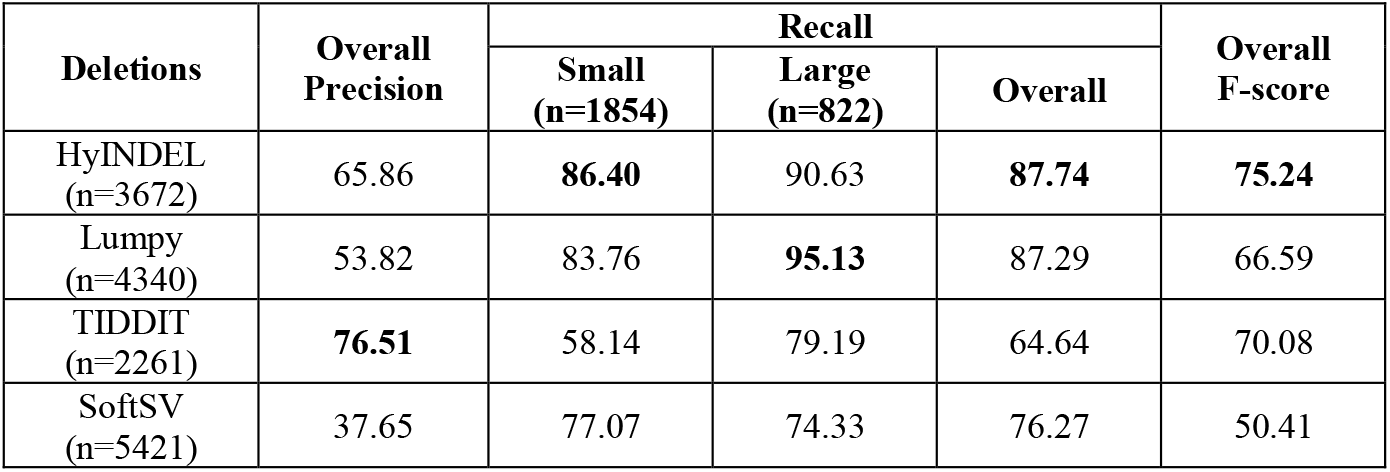
Precision, Recall and F-score metrics for predicting deletions using HyINDEL with other tools on real data (NA12878) using GIAB benchmark (n=2676)

**TABLE VI.**
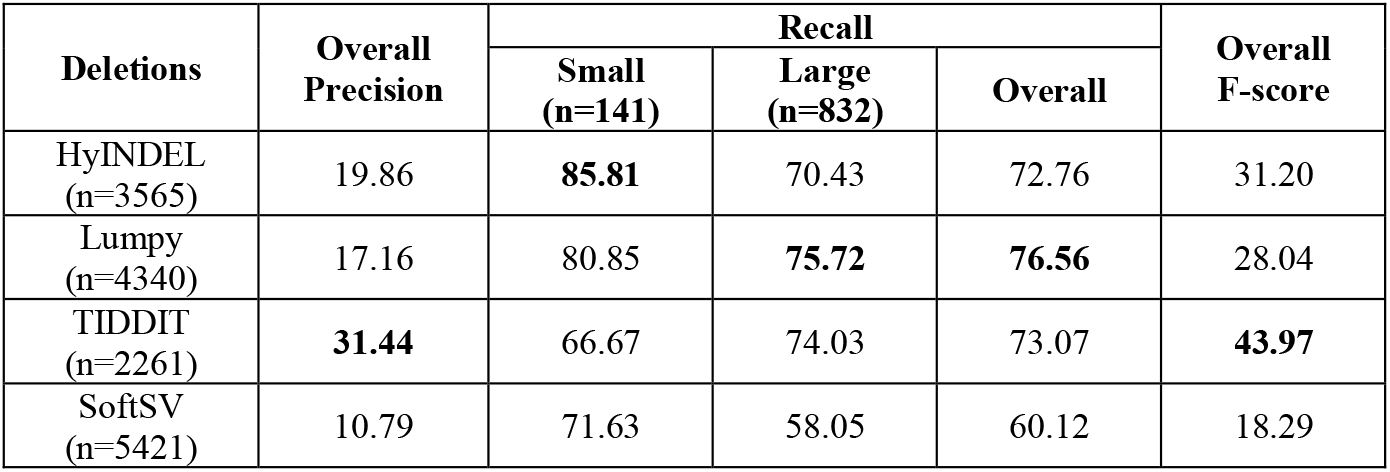
Precision, Recall and F-score metrics for predicting deletions using HyINDEL with other tools on real data (NA12878) using DGV benchmark (n=973)

It may be noted from Table V that the performance of HyINDEL is better than the other tools on GIAB benchmark dataset, as indicated by F-score values. Recall values of HyINDEL in detecting deletions is comparable with Lumpy and much higher than TIDDIT and SoftSV. Though TIDDIT exhibits highest precision (~76.5), its recall values are least suggesting that large number of deletion events are missed by it. Precision of HyINDEL is much higher than Lumpy, resulting is a higher F-score. For the DGV dataset (Table VI), HyINDEL exhibits higher recall for small deletions, while it is highest for Lumpy in case of detection of large deletions. TIDDIT again exhibits highest precision and recall values comparable with other tools, resulting in best F-score on DGV dataset. In the case of insertions (Table VII), relatively fewer predictions (870) are observed with HyINDEL compared with other tools Pamir (5820) and Popins (3204), indicating much lower false positives. On the DGV dataset, the number of true predictions by HyINDEL (65/108) is comparable with Pamir (66/108), while on the DGV dataset, HyINDEL exhibited 43/68 insertions, higher than Pamir (20/68) and comparable to Popins (53/68) on GIAB dataset, but with a significantly higher true positive rate.

**TABLE VII.** Comparison of number of true insertions by HyINDEL, Pamir and Popins with annotations in DGV and GIAB on real data (NA12878)

| Insertions | DGV (108) | GIAB (68) |
| --- | --- | --- |
| HyINDEL (n=870) | 60 | 42 |
| Pamir (n=5820) | **66** | 20 |
| Popins (n=3204) | 22 | **53** |

### C. Implementation details

HyINDEL is implemented in combination of C++ and bash. For each chromosome, it processes regions of size 10Mb (with an overlap pf 20Kb) in parallel. On a computer with two Intel E5-2640 processors having 12 cores each, that is, a total of 24 cores, and 48GB memory, it took 2h 49m for detecting insertions and deletions on real data (NA12878) sample.

## IV. Conclusions

In this work we proposed a hybrid approach for the detection of both insertions and deletions on a single platform from next generation sequencing data. Using soft-clip reads, HyINDEL is able to provide good support in accurately detecting the INDEL breakpoints compared to other methods with high precision, indicating the reliability of predictions. It is able to handle detection of both small and large INDELs and also identify the novel insertion sequence. Our analysis indicates that the performance of HyINDEL is comparable with other state-of-the-art tools on both simulated and real data. With its reliability in predictions, we propose to extend it to handle paired case-control data (e.g., tumour-normal) and multi-sample data for the detection of population specific INDELs.

